# A hierarchical cascade of sleep rhythms drive memory consolidation in humans and are disrupted in epilepsy

**DOI:** 10.1101/2025.05.06.652436

**Authors:** Anirudh Wodeyar, Dhinakaran Chinappen, Hunki Kwon, Wen Shi, Mark Richardson, Mark A Kramer, Catherine J Chu

**Author notes:** Correspondence to: Anirudh Wodeyar, Ph.D., 4^th^ Floor, Paul Henri Spaaklaan 1, Maastricht, The Netherlands 6229 EN, Catherine J. Chu, M.D., Charlotte R. Bloomberg Children’s Center Building, 3^rd^ floor 1800 Orleans Street Baltimore, MD 21287. These authors contributed equally (Mark A Kramer, Catherine J Chu).

## Abstract

The cross-regional interplay of slow oscillations, sleep spindles, and ripples during sleep is believed to support systems memory consolidation but is understudied in humans. Using a validated behavioral task and intracranial neural recordings from orbitofrontal cortex, thalamus, and hippocampus in 19 epilepsy patients, we examined the cross-regional interplay of sleep-oscillations and their role in memory consolidation. Orbitofrontal slow oscillations robustly modulate sleep rhythms both within and across regions. Most combinations of oscillation rates predict overnight memory consolidation, but hippocampal ripple rate and coupled hippocampal-orbitofrontal ripples were the strongest positive predictors of memory consolidation. In contrast, epileptic spikes coupled to sleep oscillations strongly predicted reduced memory consolidation, with the strongest negative effect observed when epileptic spikes were coupled to slow oscillations. These findings provide direct evidence of the hierarchical cascade of sleep oscillations in human memory processing and reveal how epileptic spikes disrupt this process in patients with epilepsy.

## Introduction

Sleep is a critical period for the integration and strengthening of memories. Three cardinal sleep electrophysiological rhythms - slow oscillations (0.5-2 Hz), spindles (8-15 Hz), and ripples (70-120 Hz) - have been linked to non-rapid eye movement (non-REM) sleep-dependent memory consolidation^1–4^. Available evidence suggests that slow oscillations originate primarily in the frontal cortex^5^ before propagating across widespread regions^5,6^. Regional and generalized thalamocortical sleep spindles are initiated in the thalamus, coordinated with the up-state of slow oscillations^7–9^. Ripples, a short (<100 ms) and temporally discrete oscillation in the hippocampus have been linked to offline memory replay^10,11^, and ripple-like focal events have now been detected throughout the human cortex^12–14^. The interplay of slow oscillations, spindles, and ripples across frontal cortex, thalamus and hippocampus is thought to be crucial for systems memory consolidation^15,16^, though the precise coordination of these electrophysiological events across this anatomical circuit remains poorly understood.

Previous human studies have demonstrated pairwise coupling and temporal coordination among slow oscillations, spindles, and ripples in frontal regions, hippocampus, and thalamus^1,8,12,17–21^ replicating and extending analyses from rodent models^2,3,22–24^. These studies have also shown that individual or pairwise combinations of sleep oscillations across different subsets of neocortex, hippocampus, and thalamus predict memory consolidation^2–4,11,23,25^. However, despite these important insights into sleep oscillatory coordination and memory, the optimal integration of cardinal sleep rhythms across the broader circuit connecting the frontal cortex, thalamus, and hippocampus to support human memory remains unknown.

In parallel, memory concerns are the primary cognitive symptom reported in epilepsy, a disease characterized by pathologic neuronal synchronization^26–28^. Memory symptoms in patients with epilepsy are frequent even in the absence of antiseizure medications, structural lesions, or frequent seizures. In wakefulness, epileptic spikes interfere with memory encoding^29–32^. In sleep, epileptic spikes are facilitated by slow oscillations and “hijack” or disrupt spindles and ripples^9,26,33–35^, however the impact that this disruptive interference of sleep oscillations has on sleep-dependent memory consolidation remains poorly understood.

To address these knowledge gaps, we evaluated sleep oscillations, epileptic spikes and sleep-dependent memory consolidation in a cohort of patients with drug refractory epilepsy and intracranial electrophysiological recordings simultaneously sampling that included the medial orbitofrontal cortex, thalamus, and hippocampus. We examined the spatiotemporal coordination of slow oscillations, spindles, and ripples within and across orbitofrontal cortex, thalamus, and hippocampus. We then evaluated the relationship of each oscillation and combination within and across brain regions to memory consolidation. Next, we evaluated the impact of epileptic spikes alone and co-occurring with sleep oscillations on memory consolidation. Finally, we identified the optimal predictive model of overnight memory consolidation from all combinations of events across all brain regions. Through this work, we provide evidence of a hierarchical cascade of network activity strongly modulated by frontal slow oscillations, and through which downstream ripple oscillations most effectively consolidate memory. In contrast, epileptic spikes coupled to slow oscillations at the top of this cascade disrupt this elegant process and strongly predict worsened memory consolidation. Our findings thus support and refine the general framework of systems memory consolidation and indicate how memory consolidation can be impaired in epilepsy.

## Results

### Subject Characteristics

All subjects admitted to the MGH Epilepsy Monitoring unit with stereo-electroencephalography (SEEG) electrodes sampling medial orbitofrontal cortex (OFC), thalamus, and hippocampus between 5/2021 and 11/2023 were included. Nineteen subjects (26 sampled hemispheres) met these inclusion criteria. Fourteen subjects completed the motor sequence typing task (28 hands tested), a validated behavioral assay of sleep-dependent memory consolidation^36^, before and after sleep on the night analyzed. Subject characteristics are provided in Supplementary Table 1.

### Characteristic slow oscillations, spindles, and ripples are observed in each region

Slow oscillations, sleep spindles, and ripples were detected in all channels in all three regions during all bouts of stages 2 (N2) and 3 (N3) non-rapid eye movement sleep over a 12-hour period using previously validated detectors (see *Methods*; **Figure 1**). For these analyses, sleep oscillations that co-occurred with epileptic activity in any region were excluded. We found an average rate of 5.22 / min [±0.21 / min standard error (s.e.)] for slow oscillations, 5.78 (±0.46) / min for spindles, and 3.19 (±0.21) / min for ripples. Slow oscillation and spindle rates were higher in the OFC and thalamus than in the hippocampus (comparison corrected p<0.05) while ripple rates were highest in the hippocampus (comparison corrected p<0.05). For details on rates within each region and the statistical testing, please see **Supplementary Table 1**.

**Figure 1:**
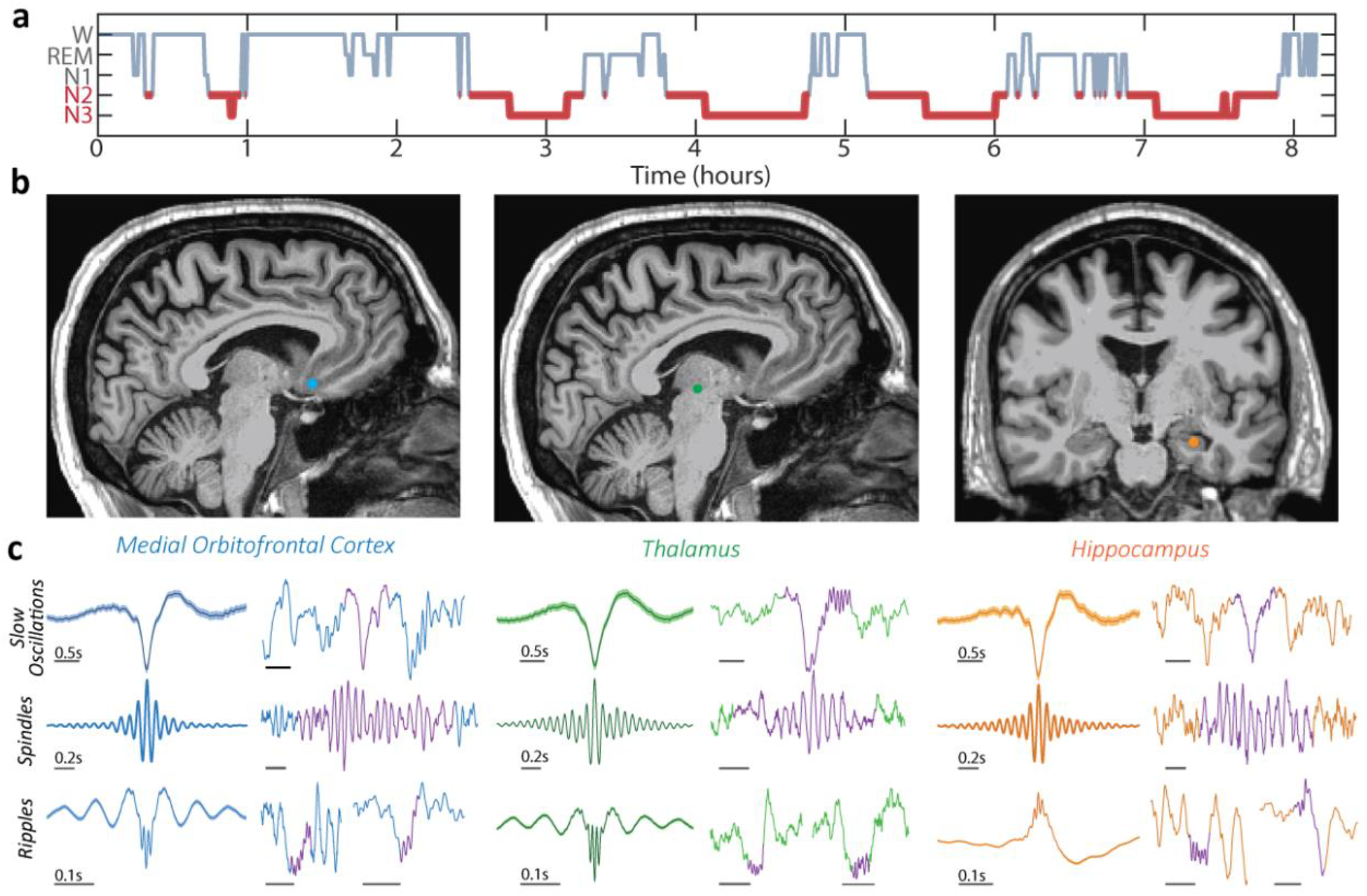
Canonical rhythms of sleep are observed in human cortex, thalamus, and hippocampus. **(a)** Example overnight hypnogram from one subject where N2 and N3 epochs were selected for analysis (red). **(b)** Example SEEG lead locations in one subject sampling OFC (blue circle), thalamus (green circle), and hippocampus (orange circle). **(c)** Example averaged (left) and individual (right, purple) slow oscillations, spindles, and ripples detected in each location. Amplitudes rescaled for visualization.

### Slow oscillations occur in the OFC first and coordinate across thalamus and hippocampus

Visual inspection of the average voltage trace across subjects demonstrates that slow oscillations are strongly temporally coordinated across regions (**Figure 2a**). A triphasic slow oscillation appeared in each region, with an initial up-state peak near -0.5 s and a second up-state peak near +0.5 s relative to the down-state trough. On average, the slow oscillation trough occurred first in the medial OFC, followed by the thalamus [15.6 ± 2.7 ms later, T(25) = 5.877, p = 3.9e-6], and then the hippocampus [35.1 ± 12.3 ms later, T(25) = 2.86, p = 0.0085]. The presence of an OFC slow oscillation predicted an up-regulation of thalamic slow oscillation occurrences by a factor of 9.51 (± 0.09, p∼0), and hippocampal slow oscillation occurrences by a factor of 5.4 (± 0.11, p, ∼0), relative to baseline rates in each region.

**Figure 2:**
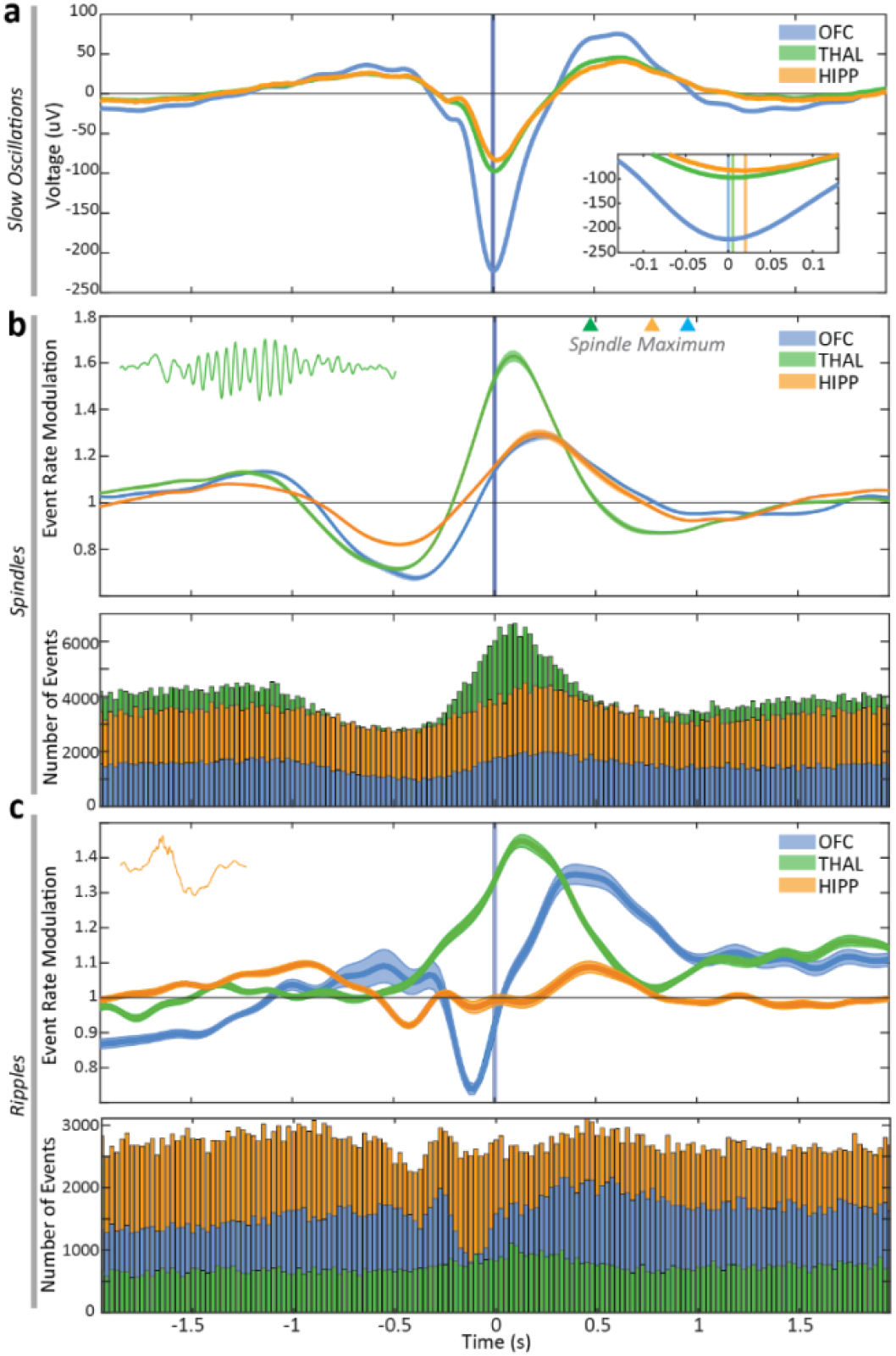
Cross-regional coupling of slow oscillations, spindles, and ripples in humans during sleep. (**a**) All detected slow oscillations temporally aligned to the medial orbitofrontal cortex (OFC, blue) slow oscillation trough. (Inset) The minima of averaged events are observed first in the OFC followed by the thalamus (green) then hippocampus (orange). (**b**) (Upper) Modulation in incidence relative to baseline spindle rates in the OFC, thalamus, and hippocampus temporally aligned to the OFC slow oscillation minima. Triangles show the spindle amplitude maxima, which coincide with the slow oscillation up-state. (Lower) Total number of spindles detected in each region (25 ms time bins). (**c**) (Upper) Modulation in incidence of ripples in the OFC, thalamus, and hippocampus temporally aligned to the OFC slow oscillation trough. (Lower) Total number of ripples detected in each region (25 ms time bins). Shaded areas indicate ±95% confidence intervals.

### Sleep spindles coordinate across distant brain regions and are modulated by OFC slow oscillations

Controlling for baseline rates, we evaluated the incidence of detected spindle onsets in each region relative to the slow oscillation trough in the OFC (see *Methods*). We found that the spindle incidence in each region was modulated by OFC slow oscillations (**Figure 2b**). Spindles in the thalamus were maximally up-regulated 0.100 s (0.025 s bins) after the OFC slow oscillation minima (by a factor of 1.66, corrected 95% confidence interval [1.63, 1.68], p≈0). Spindles in the medial orbitofrontal cortex were maximally up-regulated 0.25 s after the OFC slow oscillation minima (by a factor of 1.29, 95% CI [1.27, 1.31], p≈0). Spindles in the hippocampus were maximally upregulated 0.225 s after the OFC slow oscillation minima (by a factor of 1.26, 95% CI [1.25, 1.28], p≈0). Compared to spindle onsets, the maximum spindle amplitude in each region aligned with the slow oscillation up-state in the thalamus (at 0.475 s after the OFC slow oscillation trough), the hippocampus (0.775 s), and the OFC (0.950 s, see Figure 2B triangles and **Supplementary Figure 1**), as expected.

Sleep spindles in all three regions were also moderately - but significantly - up-regulated prior to the OFC slow oscillation trough, coinciding with the first up-state of the triphasic OFC slow oscillation (**Figure 2b**). In the OFC, the maximum up-regulation of spindle incidence occurred 1.15 s before the OFC slow oscillation minima (increase by a factor of 1.11, 95% CI [1.11, 1.12], p≈0). In the thalamus, the maximum up-regulation of spindle incidence occurred 1.25 s before the OFC slow oscillation minima (increase by a factor of 1.16, 95% CI [1.15, 1.17], p≈0). In the hippocampus, the maximum up-regulation of spindle incidence occurred 1.325 s before the OFC slow oscillation trough (by a factor of 1.09, 95% CI [1.09, 1.10], p≈0). Early spindle activity preceding the slow oscillation trough and coinciding with the first slow oscillation up-state was also observed when only isolated slow oscillations (separated by at least ±3 s from another slow oscillation) were evaluated (**Supplementary Figure 2**).

Thalamic sleep spindles had higher amplitude (mean 4.9 *μV*, s.e. 0.18) and duration (mean 1.24 s, s.e. 0.07) than hippocampal spindles (amplitude mean 3.55 *μV*, s.e. 0.14, t(25) = 8.47, p = 8.2e-9; duration mean 0.97 s, s.e. 0.07, t(25) = 6.14, p = 2e-6) and higher amplitude but not duration than OFC spindles (mean 3.65 *μV*, s.e. 0.15, t(25) = 7.45, p = 8.3e-8; mean 1.15 s, s.e. 0.09, t(25) = 1.38, p = 0.18). OFC sleep spindles had a lower mean frequency (11.9 Hz, s.e. = 0.07 Hz) than spindles detected in the hippocampus (mean 12.25 Hz, s.e. 0.08 Hz, t(25) = 3.95, p = 5.7e-4). In the thalamus, spindle frequency varied across nuclei; spindle frequencies were slower in the anterior nucleus than in the pulvinar nucleus (11.99 Hz versus 12.9 Hz, t(26) = -4.27, p = 4.6e-4). The timing of slow (9 – 12 Hz) and fast (12 - 15 Hz) spindle modulation relative to the OFC slow oscillation was similar across brain regions (**Supplementary Figure 3**).

To evaluate the impact of co-occurring thalamic spindles and OFC slow oscillations on spindle propagation, we evaluated the incidence of cortical or hippocampal spindles in the setting of thalamic spindles with and without a co-occurring OFC slow oscillation. When a thalamic spindle and orbitofrontal slow oscillation were both present, OFC spindle occurrence increased by a factor of 3.93 (95% CI [3.89, 3.98], p≈0) and hippocampal spindle occurrence increased by a factor of 3.31 (95% CI [3.29, 3.34], p≈0). In contrast, in the absence of a co-occurring OFC slow oscillation, a thalamic spindle up-regulated spindle occurrence in OFC by a factor of 1.82 (95% CI [1.78, 1.86], p≈0) and in the hippocampus by a factor of 1.61 (95% CI [1.58, 1.64], p≈0).

### Ripple oscillations coordinate across distant brain regions and are modulated by OFC slow oscillations

Sharp wave ripples, characterized by fast 80-120 Hz oscillations lasting less than 100 ms occurring alongside a 2-8 Hz wave have classically been described in the hippocampus^37^. Recently, ripple oscillations detected agnostic to sharp waves have also been reported in thalamus and cortex^12,17,38,39^. Controlling for baseline rates, we evaluated the ripple incidence in each region relative to the slow oscillation trough in the OFC (see *Methods*). We found that the ripple incidence in each region was modulated by an OFC slow oscillation (**Figure 2c**). Ripples in the thalamus were maximally up-regulated 0.125 s after the OFC minima (by a factor of 1.43, 95% CI [1.42, 1.45], p≈0). Ripples in the medial orbitofrontal cortex were maximally up-regulated 0.4 s after the OFC slow oscillation minima (by a factor of 1.34, 95% CI [1.31,1.37], p≈0). Ripples in the hippocampus were maximally up-regulated 0.475 s after the OFC slow oscillation minima (by a factor of 1.09, 95% CI [1.07, 1.10], p=2.1e-9).

Ripples in the hippocampus and OFC, but not the thalamus, were also moderately -but significantly - up-regulated prior to the OFC slow oscillation trough. The incidence of hippocampal ripples was increased 0.925 s before the slow oscillation minima (by a factor of 1.10, 95% CI [1.09, 1.10], p≈0), concordant with the first SO up-state and increase in hippocampal sleep spindles (Figure 2b). Similarly, the incidence of ripples in the OFC was increased 0.550 s before the slow oscillation minima in the OFC (by a factor of 1.07, 95% [1.03, 1.12], p=0.0011). Early ripple activity, preceding the slow oscillation trough and coinciding with the first slow oscillation peak, was also observed when only isolated slow oscillations (separated by at least ±3 s from another slow oscillation) were evaluated (**Supplementary Figure 2**).

Although ripples were detected in each location, we found that ripples in the hippocampus had higher amplitudes (13.99 uV, s.e. 1.72 uV) compared to those detected in the thalamus (3.34 uV, s.e. 0.94 uV, t(25) = 5.71, p= 5.9e-6) or orbitofrontal cortex (4.51, s.e 0.77 uV, t(25)=5.19, p=2.3e-5). We also found that ripples in the hippocampus had longer durations (43.85 ms, s.e. 0.78 ms) than those detected in the thalamus (37.2 ms, s.e. 1.0 ms, t(25)=5.08, p=3.1e-5), but no evidence of a difference in ripple durations between hippocampus and orbitofrontal cortex (41.6 ms, s.e. 1.9 ms, t(25)=1.1, p=0.28). Ripples detected in the thalamus were higher frequency oscillations (93.0 Hz, s.e. 0.3 Hz) compared to those detected in the hippocampus (85.6 Hz, s.e. 0.3 Hz, t(25) = 18.81, p=2.9e-16) or OFC (86.8 Hz, s.e. 0.53 Hz, t(25) = 11.38, p=2.2e-11; **Supplementary Figure 4**).

### Coordination of sleep oscillations across distant brain regions supports memory consolidation

To investigate whether the coupling of sleep oscillations across regions predicts overnight memory consolidation and identify the cross-regional and cross-frequency circuits that support overnight memory consolidation, we trained eligible subjects on a validated probe of sleep-dependent memory – the motor sequence typing task^44^ - before sleep and tested their performance after awakening. In total, 14 subjects completed the task (5 with bilateral sampling and bilateral hands tested, n=19 hands tested). Cross-frequency and cross-regional sleep oscillation couplings were compared to contralateral hand performance.

The rates of each sleep oscillation in each region, the rates of coupling of each sleep oscillation combination (i.e., slow oscillation-spindle, slow oscillation-ripple, spindle-ripple, slow oscillation-spindle-ripple) within each region, and the rates of coupling of each sleep oscillation combination across each region were computed (40 rates in total, see *Methods* and **Figure 3**). All analyses excluded sleep oscillations that co-occurred with epileptic activity in any region.

**Figure 3:**
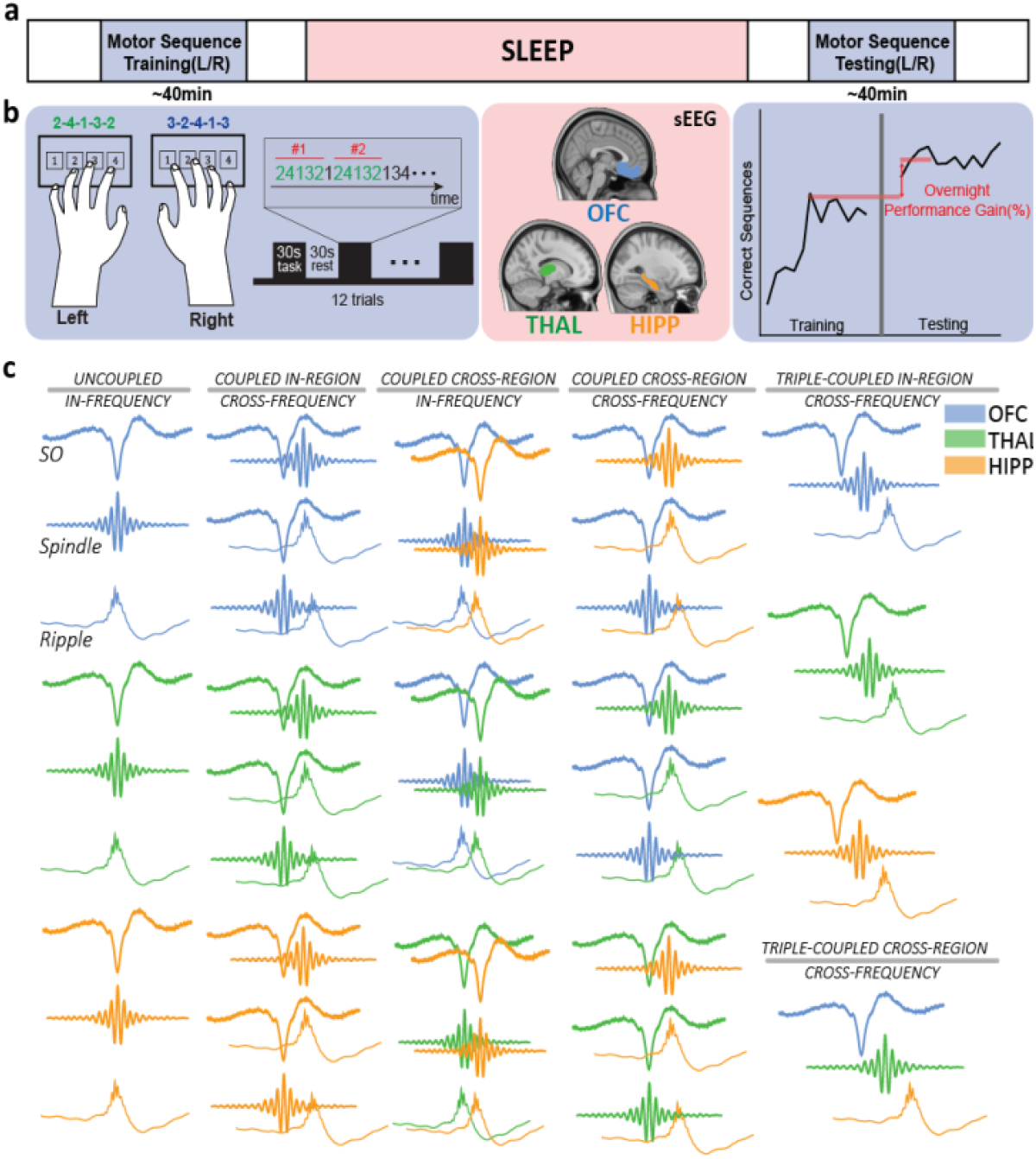
Cross-frequency and cross-regional sleep oscillations that predict memory consolidation. **(a,b)** To measure sleep-dependent memory consolidation, subjects typed a 5-digit sequence quickly and accurately for twelve 30 s trials with each hand before sleep. Neurophysiological recordings were obtained during sleep. After sleep, subjects were tested on the motor sequence task. **(c)** During stages 2 and 3 non-rapid eye movement sleep the rates of slow oscillations, sleep spindles, and ripples; all pairwise combinations of these oscillations within and across regions; and triple-coupling within and across regions were used as predictors to model overnight memory consolidation. OFC: orbitofrontal cortex; THAL: thalamus; HIPP: hippocampus.

We analyzed the correlation between each of these 40 combinations of sleep oscillations (uncoupled and coupled, within and across brain regions) and performance on the motor sequence typing task. In this univariate analysis, we found that the rate of sleep oscillations correlated positively with overnight memory consolidation for 31/40 predictors (two-sided binomial test, p = 6.8e-04, **Figure 4a**). The highest positive correlation observed was between hippocampal – OFC ripple coupling and overnight memory consolidation (r=0.49, **Figure 4c**). The strongest positive correlations all involved ripples.

**Figure 4:**
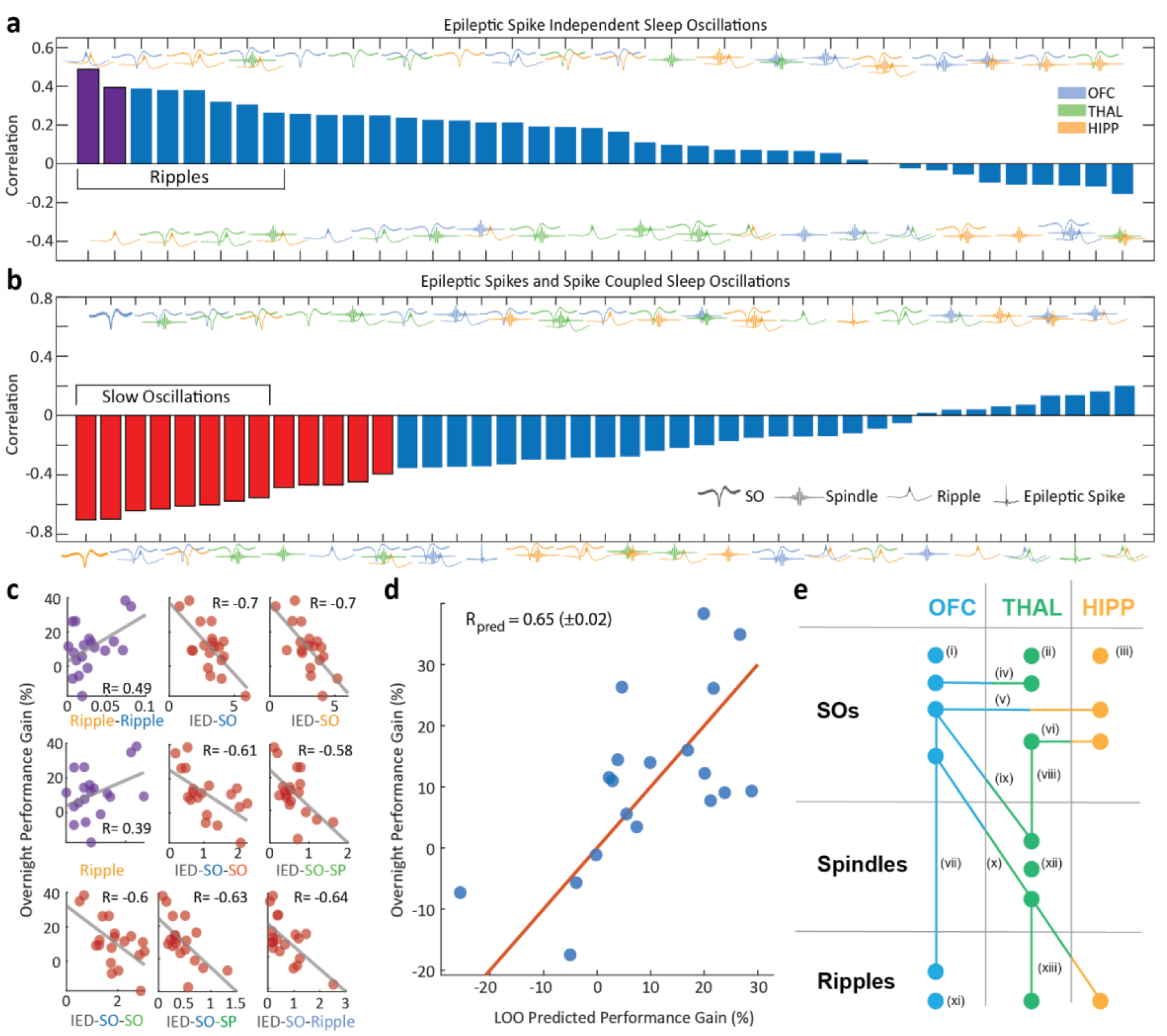
Epileptic spike-coupled sleep oscillations predict disrupted sleep-dependent memory consolidation. **(a,b)** Univariate correlations between rates of sleep oscillation predictors (a) uncoupled or (b) coupled to epileptic spikes and overnight performance gain. Purple (a) or red (b) bars represent features with significant effects across all subjects. Symbols centered on each tick mark indicate the rhythmic feature (SO, spindle, ripple, or epileptic spike) and colors indicate the brain region; see legends. **(c)** Individual scatterplots of a subset of selected predictors with overnight MST performance gain. **(d)** Prediction of overnight gain in MST performance using a partial least squares derived model versus the observed overnight performance gain. Each point indicates data from one hemisphere and the line indicates equality. R_pred_ is predictive correlation. LOO: Leave-one-out cross-validation. **(e)** Illustration of 13 predictors negatively correlated with memory consolidation. Each predictor co-occurs with an epileptic spike. SOs occurring in **(i)** OFC (r = -0.7), **(ii)** thalamus (-0.47), and **(iii)** hippocampus (r=-0.7), SOs co-occurring in **(iv)** OFC-thalamus (r=-0.6), **(v)** OFC-hippocampus (r=-0.61), and **(vi)** thalamus-hippocampus (r=-0.55). **(vii)** SOs co-occurring with ripples in OFC (r=-0.64), and **(viii)** co-occurring with spindles in thalamus (r=-0.58). **(ix)** SOs in OFC co-occurring with thalamic spindles (r=-0.63), and **(x)** triple-coupled OFC SO-thalamic spindle-hippocampal ripples (r = -0.39). **(xi)** Ripples in OFC (r=-0.47), **(xii)** spindles in thalamus (r=-0.49), and **(xiii)** co-occurring spindles and ripples in thalamus (r = -0.44).

### Co-occurrence of epileptic activity with sleep oscillations across distant brain regions disrupts memory consolidation

As all subjects included had drug refractory epilepsy, we were also able to investigate the impact of epileptic activity on memory consolidation. To do so, we detected epileptic spikes during N2 and N3 sleep using an automated epileptic spike detector (see *Methods*)^10,36,47^. We found that the rate of sleep oscillations with coupled epileptic spikes correlated negatively with overnight memory consolidation for 32/40 predictors (two-sided binomial test, p = 1.82e-04, **Figure 4b**). The highest negative correlation was between hippocampal slow oscillations co-occurring with epileptic spikes and overnight memory consolidation (r = -0.70). Across predictors, the strongest negative correlations involved epileptic spikes co-occurring with slow oscillations **(Figure 4c)**. Epileptic spike rates alone, in contrast, were only weakly correlated with memory consolidation with inconsistent relationships seen across regions (OFC: r=-0.34; THAL: r=0.14; HIPP: r=-0.12).

### Optimal predictors of sleep-dependent memory consolidation in epilepsy

To identify the optimal predictors for memory consolidation from the 83 assessed individual and coordinated sleep oscillation rates, we first identified predictors significantly coupled to motor sequence task performance for all subjects in a leave-one-out predictor selection procedure (see *Methods*; purple and red bars in Figures 3a,b respectively). Of the 83 original predictors, a subset of 15 predictors were selected. This subset of predictors included two positive predictors of memory consolidation: the rate of ripples co-occurring in the hippocampus and the OFC (without epileptic spikes) (r=0.49) and the rate of hippocampal ripples (r=0.39). The remaining 13 predictors negatively correlated with memory consolidation (Figure 4e). Each of these negative predictors included sleep rhythms with co-occurring epileptic spikes and the majority (9 of 13) included slow oscillations. Notably epileptic spike rate alone was not selected by our procedure, i.e., epileptic spike rate alone was not significantly coupled to motor sequence task performance in all subjects.

Applying partial least squares regression (see *Methods*) to these 15 predictors, we found the top performing model explained 42.25% of the variance in overnight memory consolidation (leave-one-out cross-validation predictive R = 0.65, s.e. 0.01, p≈0). We find that hippocampal-OFC coupled ripple rate is the strongest single positive predictor and that the rate of epileptic spikes coupled to sleep oscillations in the OFC and hippocampus is the single strongest negative predictor of sleep-dependent memory consolidation in this model. We conclude that coordinated sleep oscillations across the OFC, thalamus, and hippocampus support memory consolidation and that epileptic spikes coupled to these rhythms disrupts this fundamental memory process.

### Conceptual model of the hierarchical coordination of sleep oscillations supporting memory consolidation

Our findings support a model in which slow oscillations in the orbitofrontal cortex serve as primary organizers of network oscillatory activity during stages 2 and 3 non-rapid eye movement sleep. Slow oscillations travel rapidly from the frontal cortex to the thalamus and hippocampus. In the thalamus, the slow oscillation facilitates thalamocortical and/or thalamo-hippocampal spindles during the down-to-upstate transition. The slow oscillation up-state also comodulates and coordinates hippocampal and orbitofrontal ripples. This orchestrated cascade of activity across the network is graphically represented in Figure 5a.

**Figure 5:**
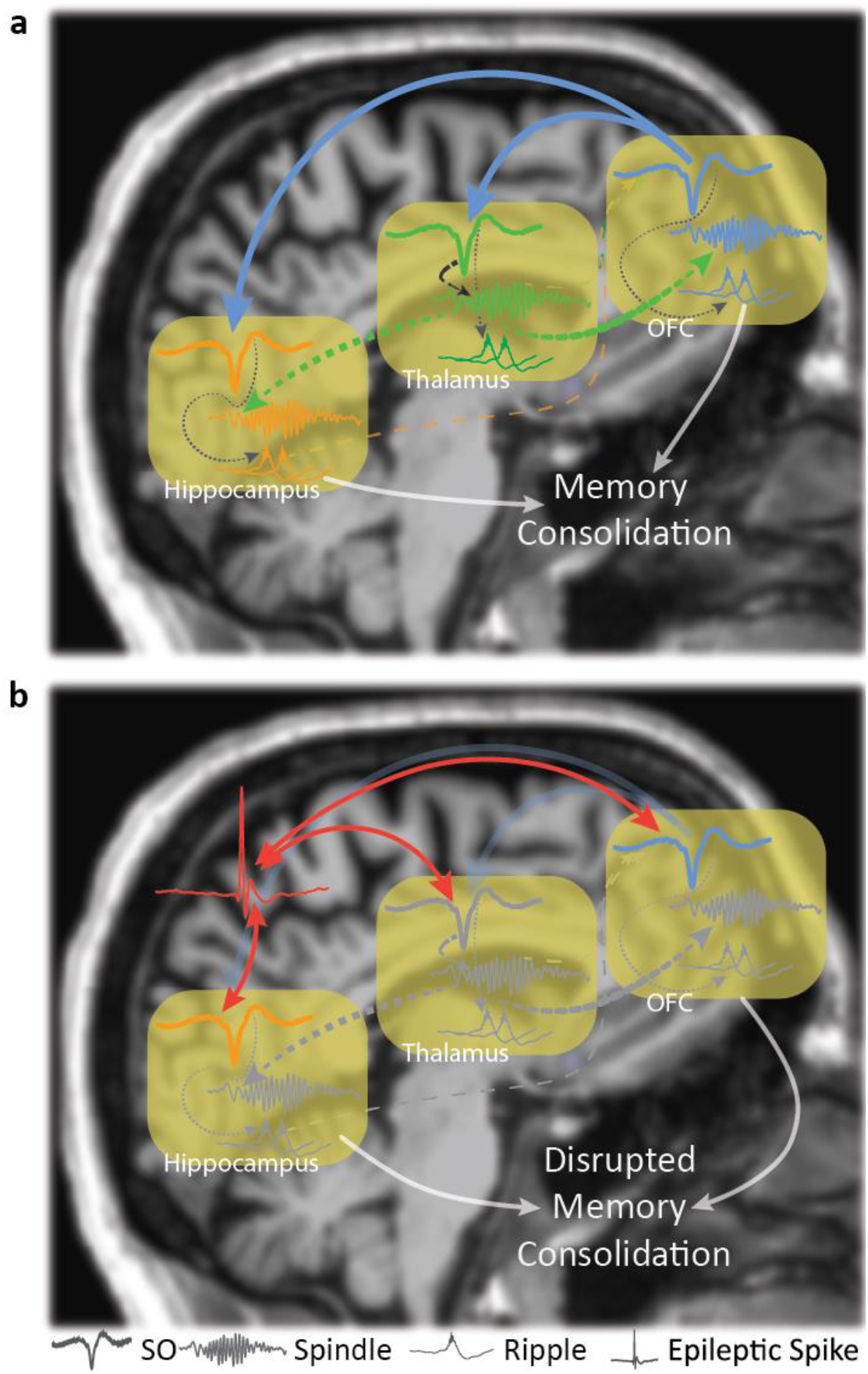
The hierarchical cascade of sleep oscillations underlying memory consolidation and their disruption in epilepsy. **(a)** Slow oscillations in the orbitofrontal cortex travel rapidly (blue arrows) to the thalamus and hippocampus. Slow oscillations facilitat thalamocortical (green/blue) and thalamohippocampal (green/orange) spindles on the down-to-upstate transition (black arrow) . Slow oscillation facilitate ripples during the up-states. Ripples coordinated across the hippocampal and frontal cortex maximally facilitate overnight memory consolidation (white arrows). **(b)** Epileptic spikes coupled to slow oscillations (red arrows) disrupt memory consolidation, potentially by corrupting the information transfer meant to flow through this hierarchical cascade of sleep rhythms.

### Conceptual model of sleep-dependent memory impairment in epilepsy

Our findings suggest a model where memory impairment in epilepsy results from the functional disruption of coordinated sleep oscillations by co-occurring epileptic spikes. Although slow oscillations normally orchestrate memory consolidation through spindle and ripple facilitation, in epilepsy, slow oscillations can also facilitate epileptic spikes on the initial up-state^9,40^. As observed in rodent models, when a spike is coupled to a slow oscillation, it effectively hijacks^26,33^ this foundational oscillatory event and disrupts the downstream physiological cascade. Notably, we find that the rates of epileptic spikes, coupled to slow oscillations were the strongest negative predictors of overnight memory consolidation. This disrupted oscillatory cascade is illustrated in Figure 5b.

## Discussion

In this manuscript, we show the hierarchical coordination of sleep oscillations that supports human memory consolidation and how epileptic spikes hijack this process and disrupt memory in epilepsy. We show that orbitofrontal slow oscillations modulate a cross-regional cascade of rhythms in which ultimately, the downstream rate of hippocampal ripples - coupled to orbitofrontal ripples or alone - best predict successful memory consolidation. Critically, the coupling of epileptic spikes to sleep oscillations, especially slow oscillations, strongly disrupts overnight memory consolidation.

The sequential pattern we observed, whereby orbitofrontal slow oscillations modulate thalamic spindles and hippocampal ripples, supports the established framework proposed for systems memory consolidation^15,16,41^. In this framework, slow oscillations create windows of excitability, spindles facilitate plasticity, and ripples reactivate or “replay” action potential sequences to enable cross-regional synaptic strengthening for long term memory storage. Our data are consistent with observations that slow oscillations initiate in the frontal cortex^5,6^ to organize widespread activation across the thalamus and hippocampus. Further, our finding that coupled hippocampal and orbitofrontal ripples are strong positive predictors of memory consolidation is consistent with prior observations that hippocampal replay is directed towards the orbitofrontal cortex for integration and long-term storage^1,12,17^. Additionally, we found a modest up-regulation of thalamic spindles and hippocampal ripples preceding the first slow oscillation up-state, also noted by others^17,42,43^. As this could indicate a preparatory or bi-directional information flow, these subtle temporal dynamics require further investigation, potentially through interventional studies.

A key finding of our study is that the negative impact of epileptic spikes on memory consolidation is not primarily linked to the overall burden of epileptic spikes^32,33,44,45^, but rate of epileptic spikes coupled to critical sleep oscillations. In past work, interactions between epileptic spikes and sleep oscillations have been recognized as a significant factor in the memory impairment seen in epilepsy^9,32,33,40,46–48^. We find that epileptic spikes, coupled with slow oscillations in any of the regions studied, were strong negative predictors of overnight memory consolidation. As slow oscillations orchestrate the cross-regional slow-oscillation-spindle-ripple cascade, spikes interfering with the foundation of this hierarchy could prevent^9,31,46^ or distort^46,49,50^ the intended information transfer typically supported by spindles and ripples. These results suggest a potential therapeutic direction to improve memory function in epilepsy is to decouple spikes from slow oscillations.

Ripples and ripple-associated replay have classically been associated with the unique architecture and function of the hippocampus^37^. However, evidence now exists that the neocortex and the thalamus produce discrete bursts of fast oscillations that approximate hippocampal ripple morphologies and, at least in the neocortex, can show similar replay^12,13,39^. Indeed, these ripple-like events also couple strongly with slow oscillations and spindles^12,51^, although the impact on memory in humans is less certain. Notably, we found that orbitofrontal and hippocampal ripples correlated with improved memory consolidation while thalamic ripples did not, suggesting that thalamic ripples may have a different function than overnight memory consolidation. In contrast, our work supports prior finding in rodent models that prefrontal cortex and hippocampal ripples potentially facilitate the targeted reactivation and strengthening of specific memory engrams^23,52^.

Spike-independent triple-coupling of slow oscillations, spindles, and ripples was not selected in our analysis to predict memory consolidation. This result suggests that the hierarchical cascade of sleep oscillations may primarily function, with respect to systems memory consolidation, to facilitate ripples, which do strongly predict memory consolidation. In contrast, we found triple-coupling that co-occurred with epileptic spikes more strongly impaired memory than spikes coupled to hippocampal ripples alone. This finding suggests that when present in the full cascade of triple coupled rhythms, epileptic spikes may expand, scramble, or distort replay and memory processes more so than an epileptic spike can unleash on a focal ripple alone^26,29,30,32,53–56^.

The motor sequence task (MST) used in this study engages procedural memory. The MST corresponds to functional activation of the hippocampus and frontal cortex^57,58^ and benefits from sleep-dependent consolidation, potentially via ripple-mediated replay^2,59^. Indeed, we observed that hippocampal-orbitofrontal cortex ripple coupling positively predicted overnight MST performance gain, possibly reflecting the consolidation of spatial or sequential aspects of this task. While MST served as a robust assay for the network dynamics analyzed here, future work should investigate whether the observed coordination and disruption patterns also generalize to other forms of memory, such as declarative memory and episodic memory.

Our findings are based on recordings from patients with epilepsy. While these data enable a unique opportunity to observe widespread brain dynamics and probe the impact of epileptic disruptions on sleep oscillations and memory, patients with epilepsy also have a range of neurophysiological abnormalities. Despite the disparate epilepsy etiologies evaluated, we saw consistent relationships between sleep oscillations and memory, as has previously been reported in both patients with epilepsy and controls^33,34^. Further, the predictive model we report here requires confirmation in larger, independent cohorts. Future work exploring coordination across different sleep stages, early versus late sleep, and incorporating the contributions of neural activity within other brain regions linked to procedural memory, such as the striatum^59^ will provide a more complete picture. Investigating the precise timing of epileptic spikes within the slow oscillation cycle and their relationship to subsequent spindle and ripple activities could offer deeper mechanistic insights^40,55^.

In conclusion, our findings reveal a complex interplay of sleep oscillations across the orbitofrontal cortex, thalamus, and hippocampus that supports sleep-dependent memory consolidation. The disruption to memory caused by epileptic spikes coupled to this oscillatory cascade underlines the role of these oscillations in memory and identifies a mechanism by which epileptic spikes disrupt cognition in patients with epilepsy. These results deepen our understanding of systems-level memory consolidation and identify mechanisms to target to advance treatments for memory dysfunction in epilepsy.

## Methods

### Subject data collection

We prospectively recruited patients undergoing clinical evaluation for drug-resistant epilepsy at Massachusetts General Hospital who had simultaneous stereo-electroencephalography (sEEG) recordings from the medial orbitofrontal cortex (OFC), thalamus, and hippocampus, along with scalp EEG. The orbitofrontal cortex was selected due to its dense recurrent connectivity with the hippocampus^60^ and thalamus^61–63^, as well as its role in processes of memory^64^. All patients provided informed consent, and the study was approved by the Massachusetts General Hospital Institutional Review Board.

Electrode localization was performed using pre- and post-operative high-resolution MRI scans co-registered with post-implantation CT scans. We first visually assessed the gross electrode positions in the medial orbitofrontal cortex, thalamus, and hippocampus of sEEG electrodes for which these areas were the intended targets during surgical placement. To refine localization, we analyzed the electrophysiological signals along each electrode shaft, identifying transitions into white matter by noting sharp reductions in signal amplitude. This two-step approach allowed for improved anatomical localization of recording sites. Specific thalamic and hippocampal subregions sampled for each subject are detailed in Supplementary Table 2.

### EEG data collection, pre-processing, and channel selection

Intracranial EEG data were collected using the clinical Natus Quantum system (Natus Neurology Inc., Middleton, WI, USA). Depth electrodes (PMT Depthalon depth platinum electrodes with 3.5 mm spacing, 2 mm contacts, and 0.8 mm diameter; or AdTech depth platinum electrodes with 5-8 mm spacing, 2.41 mm contact size, and 1.12 mm diameter) were placed in the regions of clinical interest and sampled at 1024 Hz. Using only intracranial electrophysiology, we sleep staged based on a validated algorithm that was intended for use in patients with epilepsy^65^. This algorithm has shown 85% accuracy at detecting non-rapid eye movement sleep stages using slow oscillation and spindle activity across electrodes. We analyzed only non-REM stage two and stage three sleep segments (N2 and N3) for analysis (mean 276 mins, minimum 146.5 mins, maximum 438.5 mins). All N2 and N3 sleep segments over one night were concatenated for analysis.

For slow oscillation detections, we used all electrodes from the medial OFC, thalamus, and hippocampus with a midline, far-field non-cephalic reference over the second cortical spinous process. For spindle, ripple, and epileptic spike detections, we examined adjacent medial OFC, thalamic, and hippocampal depth electrodes in a bipolar reference. We applied bipolar referencing for improved spatial and temporal resolution for these higher frequency, spatially localized oscillations.

### Epileptic spike detection

To detect epileptic spikes, we used a previously developed method^9,66^. First, we bandpass filtered the data both forwards and backwards using a finite impulse response (FIR) filter between 25 to 80 Hz (MATLAB function *filtfilt*). We then applied a Hilbert transform, calculated the analytic signal, and the amplitude envelope of this signal. For each voltage signal, the moments when the amplitude envelope exceeded three times the mean amplitude were identified as candidate spikes. To ensure the candidate spikes were not due to gamma-band oscillatory bursts or spindles, we also calculated the regularity of oscillations in an interval (±0.25 s for gamma, ±0.3125 s for spindles) around each candidate spike. To assess the regularity of the signal in this interval, we computed the Fano factor^67^ estimated by: (i) detrending the interval of unfiltered data at each candidate spike, (ii) identifying peaks and troughs after limiting maximum delays between extrema (between peaks or between troughs) to 1/50 s for gamma-band oscillatory bursts or 1/20 s for spindles, (iii) calculating the inter-peak and inter-trough intervals, and (iv) estimating the ratio of the variance of the inter-peak and inter-trough intervals to the mean of the inter-peak and inter-trough intervals. We then removed candidate spikes if the maximum deviation (calculated by rectifying the data and identifying the maximum voltage) of the unfiltered data was below three times the mean amplitude or if the Fano factor was less than 2.5 for gamma-band oscillatory bursts or less than 5 for spindles. We additionally required that the candidate spike peak was at least twice the value of the nearest neighboring peak. Of the resulting spikes, those detected within 20 ms of one another were merged into a single spike detection. We visually examined individual detections and the averaged spike waveform for each subject to confirm that the method accurately detected epileptic spikes in cortical, hippocampal, and thalamic recordings. We show example averaged spike waveforms and the time-frequency spectrogram in Supplementary Figure 4.

### Slow oscillation detection

We applied a method consistent with prior publications^9,15,20,68^ to capture the 0.75 to 1 Hz range of slow oscillations. Briefly, slow oscillations were detected by: (i) filtering the data (both backward and forwards) into the 0.4 Hz to 4 Hz band using a FIR filter (4092 order); (ii) identifying all negative to positive zero-crossings and the time interval *t* between subsequent negative to positive zero-crossings; (iii) retaining all time intervals of duration 0.75 s ≤ t ≤ 3 s (corresponding to oscillations between ∼0.4 Hz to 1.33 Hz, due to filtering) and identifying the negative peak and peak-to-peak amplitude; (iv) omitting intervals in the bottom 75th percentile of peak-to-peak amplitude and detections with peak greater than 1000 µV or trough less than -1000 µV; (v) retaining any remaining intervals as slow oscillations. When estimating the cross-correlation histograms, we omitted slow oscillations that co-occurred with an epileptic spike within 0.5 s prior to the downstate and 1 s after the downstate (based on results from Wodeyar et al., 2024^9^).

### Spindle detection

We applied an existing spindle detector with robustness to epileptic spikes and sharp transients^33^. The spindle detector estimates a latent state - the probability of a spindle - using three features: the Fano factor (estimated for data FIR filtered between 3 – 25 Hz), normalized power in the spindle band (9 – 15 Hz), and normalized power in the theta band (4 – 8 Hz). Distributions of expected values for these parameters were determined using manually detected spindles in the scalp EEG of subjects with sleep-activated spikes. The spindle detector estimates the probability of a spindle in 0.5 s intervals (0.4 s overlap). We detected a spindle when the probability crossed 0.95, chosen by optimizing the sensitivity and specificity of the detector. We retained spindles with a minimum duration of 0.5 s, a maximum duration of 5 s, and separated by at least 0.5 s; spindles within 0.5 s of one another were merged into a single spindle detection. We visually confirmed that the method accurately detected sleep spindles in OFC, hippocampal, and thalamic recordings and omitted all spindles that had a co-occurring spike within the spindle duration for cross-correlation histogram analysis.

We estimated spindle frequencies following a method similar to that proposed in Kwon et al. (2023)^69^. To do so, we: (i) ensured each spindle reached duration 10 s; (ii) applied a Hanning taper; (iii) computed the spectrum; (iv) removed from the spectrum the 1/f-like background estimated between 4-50 Hz; (v) determined the frequency of maximum power between 9–15 Hz in the resulting spectrum. We repeated this process for all detected spindles.

### Ripple detection

We detected ripples using a validated method adapted from Staba et al., (2002)^70^ and applied in several studies^18,71^ . For a given bipolar signal, we: (i) bandpass filtered the signal between 70 to 150 Hz; (ii) applied the Hilbert transform to compute the ripple-band envelope; (iii) estimated the root-mean-squared amplitude of the ripple-band envelope and set the threshold for ripple detection at the 97.5th percentile, similar to that used in other studies^20,71,72^; (iv) retained ripples that exceeded the 97.5th envelope threshold for at least 6 ms, produced at least 6 voltage peaks in both the raw and the bandpass filtered signal, and possessed a ripple-band envelope >3 SDs above the mean for all values in a 200 ms interval centered on the ripple-band envelope peak. For the cross-correlation histogram analysis, we omitted ripples that co-occurred with a spike within 0.5 s. Ripple spectrograms computed using a multitaper spectral estimate with 2 Hz bins, standardized within subject, and averaged across subjects are shown in Supplementary Figure 5.

### Temporal Coupling with Cross-Correlation Histograms

We estimate cross-regional and cross-frequency relationships using event-based covariance statistics, specifically the normalized cross-correlation histogram^9^. In this approach, we use event detections to estimate relationships between regions and oscillations, thus minimizing bias from interictal activity in the original SEEG data. For each slow oscillation detection, we assigned the event time to the downstate minimum. For each spindle detection, we assigned the event time to either the spindle start time or the maximum amplitude depending on the analysis performed. For each ripple detection, we assigned the event time to the moment of maximum amplitude. We compute the cross-correlation histogram between two rhythms (rhyhtm_1_ and rhythm_2_) in two regions (region_1_ and region_2_) as follows: (i) create binary time series for each region, marking the event times of rhyhtm_1_ in region_1_ and rhyhtm_2_ in region_2_; (ii) determine the time *t*_≤_ of each event ≤ from the binary time series of rhyhtm_1_ in region_1_; (iii) compute the delays between all events in the binary time series from rhyhtm_2_ in region_2_ within ≤ s of each *t*_≤_ . These delays for all channel pairs in the two regions within each subject were computed.

To estimate the population-level distribution of event incidence around slow oscillations, we employed the following bootstrapping approach: (i) For every iteration of the bootstrap, we sampled randomly with replacement from all hemispheres of all subjects to select 26 hemispheres and concatenated all delays relative to downstate event detections from each selected hemisphere. (ii) We generated a smoothed estimate of the histogram of the resampled delays using 75 ms kernel smoothing (MATLAB *fitdist*). (iii) We calculated the histogram in 25 ms intervals from -3 s to +3 s relative to a slow oscillation downstate minimum. We repeated the entire procedure 100 times to generate 100 resampled histograms. We then calculated the up or down-regulation of event occurrences by dividing the histogram between [-2,2] s by the average of the histogram in the -2.8 to -2 s and 2 to 2.8 s intervals (avoiding edge effects from normalization). The mean and standard error of the normalized histogram were estimated. We repeated the procedure above to assess temporal relationships between slow oscillations across regions, between OFC slow oscillations and spindles across regions, and between OFC slow oscillations and ripples across regions. Separately, we calculated the total number of events using unnormalized histograms with 25 ms time bins, summing across all subjects.

### Finger tapping motor sequence task

For the memory task, we used a validated probe of sleep dependent memory consolidation, the finger tapping Motor Sequence Task (MST)^36^. This task captures sleep-dependent memory consolidation in healthy adults, where performance improves after sleep compared to an equal period of wakefulness. Subjects were asked to place their fingers on four numerically labeled keys on a standard number pad. They were then instructed to repeatedly type a 5-digit sequence (e.g., 4-1-3-2-4) as quickly and accurately as possible during twelve 30 s trials separated by 30 s rest periods (Figure 3a). To minimize the involvement of working memory, the sequence was displayed on a monitor during the typing trials. Subjects were trained on separate sequences with their left and right hands before sleep, with a 10 min break between left- and right-hand training sessions. Following sleep, subjects were tested on the same sequences on the left and right hand respectively with an intervening 10 min break.

MST performance in the hospital setting can yield performance outliers due to, for example, inattention, task interruption, or misplacement of the fingers on the keys. To identify and exclude outliers in each trial, we followed the procedure in Manoach et al. (2004)^73^. For each subject and each hand, we fit an exponential model to the learning curve during training and included a constant offset to the exponential model during the post-sleep testing. If performance on a trial exceeded two standard deviations below the model fit, it was excluded as an outlier. To quantify sleep-dependent memory consolidation for each hand, we calculated the percentage difference between the mean of the last three trials in a training session and the mean of the first three non-outlier sequences in the testing session, following previously published procedures^36^.

### Sleep Oscillation Metrics

We detected slow oscillations, spindles, and ripples in all three regions analyzed and used these detections to estimate average rates (the number of detections divided by the total recording duration in minutes) within each region for each subject. We also measured the co-occurrence rates of slow oscillations, spindles, and ripples across regions, as well as rates of slow oscillation and spindle co-occurrence, slow oscillation and ripple co-occurrence, and spindle and ripple co-occurrence within and across regions. Finally, we calculated the triple-coupling between slow oscillations, spindles, and ripples both within region and across regions. For a visual representation of all 40 occurrences and co-occurrences computed, see Figure 3c.

We define co-occurrence as follows. For slow oscillations, co-occurrence denotes events that occur within ±1 s of the downstate trough; for spindles, co-occurrence denotes events that occur within ±0.5 s of the spindle maximum; for ripples, co-occurrence denotes events that occur within ±0.1 s of the ripple maximum. When calculating co-occurrence between a lower frequency oscillation and higher frequency oscillation, the lower frequency oscillation temporal interval was used to determine event co-occurrence.

To assess the influence of epileptic spikes on overnight memory consolidation, we partitioned event rates into two categories. In the first category, we selected sleep oscillation events without a co-occurring epileptic spike in any region. To do so, we omitted: slow oscillations that co-occurred with an epileptic spike in a [-0.5, 1] s interval around the downstate trough, spindles with an epileptic spike that co-occurred during the spindle duration, and ripples that co-occurred with an epileptic spike in a [-0.5, 0.5] s interval around the peak ripple amplitude. In the second category, we analyzed sleep oscillation events with a co-occurring epileptic spike. To do so, we included sleep oscillation events for which an epileptic spike occurred within the intervals defined for the first category. Finally, we calculated epileptic spike rates within each region as the average spike rate of all channels in a region. In total, we obtained 40 features in the first category of sleep oscillation events without epileptic spikes, 40 features in the second category of sleep oscillation events with epileptic spikes, and 3 epileptic spike rate features (one in each region), yielding a total of 83 features. We estimated univariate Pearson correlations between all features and the overnight memory consolidation.

### Partial Least Squares Regression

Given the large number of features (k=83), the relatively small number of samples (n=19 subjects), and the high level of correlation among the features, we applied the partial least squares (PLS) regression technique^74^ to characterize the relationship between oscillatory coupling and memory consolidation. PLS, a multivariate statistical analysis technique robust to highly correlated predictors, has been successfully applied to assess relationships between oscillatory coupling and behavior^75,76^.

PLS is analogous to principal components analysis in that components are created to optimally predict a response (dependent variable). Here the response *Y* is the overnight performance gain and the predictors *X*are the 83 sleep oscillation and epileptic spike features. After standardizing the predictor *X* and response *Y* variables, we apply the *plsregress* function in MATLAB which implements the SIMPLS ^74^ algorithm. SIMPLS iteratively retrieves the first left singular vector (i.e., the vector with the largest eigenvalue) from the decomposition of the matrix *X′Y*, removes the subspace associated with that singular vector from *X′Y*, and repeats the process. Subsequently, a small number of singular vectors are used instead of the many original predictors to generate a set of parameter estimates to predict the response *Y*.

To improve performance of the PLS model, we implemented predictor selection^77^ as a first step. To select candidate predictors, we computed the univariate correlation between each feature and the response and retained a feature as a predictor in the PLS model if the p-value of the univariate correlation (calculated using a t-statistic with MATLAB’s corrcoef function) was p<0.1. We chose the threshold of p<0.1 since univariate correlations do not account for the unique variance explained by each feature in a multivariate model. We applied this candidate predictor selection procedure for each leave-one-out subset of samples. The leave-one-out procedure calculated the correlation and p-value between each candidate predictor and the response over 18 samples and repeated this process for each leave-one-out subset. This candidate predictor selection procedure resulted in *n* = 19 sets of candidate predictors, one set for each left out sample. To identify the predictors that occurred consistently across these *n* sets, we applied a binomial test (*n* = 19, assuming a null probability of 0.10 given the threshold p-value selected) to the *n* sets of candidate predictors. Doing so resulted in selection of 15 of 83 predictors (see *Results*) for analysis. To assess the predictive strength of the PLS model with these selected predictors, we performed leave-one-out cross-validation, training the PLS model on the selected predictors from each subset of *n* − 1 subjects. We applied the PLS model with 4 singular vectors, though other values for this hyperparameter yielded similar predictive ability for the model. We did not assess performance of the PLS model on any training data as the model is intentionally overfit to these data. We bootstrapped the response over the n PLS model predictions, (i.e., we sampled with replacement from all pairs of predictions and response and re-calculated the correlation for each bootstrapped dataset) to estimate the standard deviation for the predictive Pearson correlation.

### Statistical Analysis

For the cross-correlation histograms, we tested whether event probabilities were significantly up-regulated or down-regulated relative to the baseline (value of 1). To account for multiple comparisons across the 2 s interval, we employed Bonferroni correction on the confidence intervals – a conservative approach to inference when there are multiple comparisons^78^. We report the mean and Bonferroni-corrected 95% confidence intervals for all instances of up- or down-regulation of event probabilities. P-values were generated as a dual of the confidence interval, as is appropriate for bootstrapping-derived inference^79^ with the null hypothesis assumed to be a value of 1 which was equivalent to no modulation of the rates of events. Rhythm characteristics were tested using paired t-tests and were determined significant after Bonferroni correction, when appropriate. We used a Binomial test to assess significance of the number of positive and negative correlations of events rates with overnight performance gain assuming the null hypothesis of an equal chance of positive or negative correlations. We used partial least squares regression with predictor selection to predict memory consolidation and reported the correlation coefficient (R) between predicted and observed memory consolidation performance gain with bootstrapped confidence interval (see ***Partial Least Squares Regression***). All statistical tests were two-tailed with a significance level of ≤ = 0≤. Analyses were performed using MATLAB (version R2019b, The MathWorks Inc., Natick, MA).

### Resource availability

#### Software and Algorithms

|  |  |  |
| --- | --- | --- |
| MATLAB | MathWorks |  |
| Spindle Detection | Kramer et al.<sup>33</sup> | https://github.com/Mark-Kramer/Spindle-Detector-Method |
| Original Code | This paper | https://github.com/wodeyara/TripleCoupling |

#### Lead contact

Further information and requests for resources should be directed to and will be fulfilled by the corresponding authors.

#### Materials availability

No materials were generated in this study.

## Acknowledgements

We thank Erin D. Berja, Katherine G. Walsh Jonathan F. Huang, Elizabeth A. Kinard and Skyler K. Goodman for their support in data collection. The authors were supported by NIH NINDS R01NS115868 and the Epilepsy Foundation New England Blue Skies Award.

## Author Contributions

Conceptualization, A.W., C.J.C., M.A.K.; Investigation, D.K, H.K., W.S., A.W.; Software, A.W., M.A.K, H.K.; Data Curation, D.K, W.S., A.W.; Formal Analysis: A.W.; Writing – Original Draft: A.W.; Writing – review and editing, A.W, C.J.C, M.A.K.; Supervision: C.J.C, M.A.K.

## Data and Code availability

Data needed to reproduce figures from this paper and code to reproduce analyses will be placed in a publicly available repository prior to publication. The raw data reported in this study cannot be deposited in a public repository because of privacy concerns. To request access, contact the lead author.

