## Supplementary Information for "A hierarchical cascade of sleep rhythms drive memory consolidation in humans and are disrupted in epilepsy"

Supplementary Table 1: Sleep Oscillation Event Rates

|  | Slow Oscillation | Spindle | Ripple |
| --- | --- | --- | --- |
| OFC | 5.1 (0.26) | 5.83 (0.89) | 3.85 (0.25) |
| Thalamus | 6.8 (0.2) | 6.63 (0.61) | 1.51 (0.11) |
| Hippocampus | 3.76 (0.31) | 3.2 (0.57) | 4.47 (0.41) |

T(25)SO:OFC-THAL = -10.78, p = 6.9e-11; T(25)SO:OFC-HIPP = 6.64, p = 5.9e-7; T(25)SO:THAL-HIPP = 11.39, p = 2.1e-11

T(25)SP:OFC-THAL = -1.23, p = 0.23; T(25)SP:OFC-HIPP = 4.33, p = 2.1e-4; T(25)SP:THAL-HIPP = 7.46, p = 8.1e-8

T(25)RIPP:OFC-THAL = 7.58, p = 6.2e-8; T(25)RIPP:OFC-HIPP = -1.86, p = 0.079;
T(25)RIPP:THAL-HIPP = -7.36, p = 1.1e-7

Legend: SO = slow oscillation; OFC = Orbitofrontal Cortex; THAL = Thalamus; HIPP = Hippocampus; SP = Spindle; RIPP = Ripple.

Supplementary Table 2: Subject Characteristics

Part A: Summary

| SEX | Age | N2 & N3 (min) | RIGHT THAL | LEFT THAL | RIGHT HIPP | LEFT HIPP | MST |
| --- | --- | --- | --- | --- | --- | --- | --- |
| 8 M, 11 F | Mean 33 (Range 14 - 65) | 276.13 (34.88) | ANT (12); CM (5);  PUL (5) | ANT (13); CM (5); PUL (5) | aHC (10), pHC (10) | aHC (14), pHC (8) | L (11), R(8) |

Part B: Subject-Specific Details

| ID | AGE | SEX | N2 & N3 (min) | RIGHT THAL | LEFT THAL | RIGHT HIPP | LEFT HIPP | ASM | ETIOLOGY | MST |
| --- | --- | --- | --- | --- | --- | --- | --- | --- | --- | --- |
| 1 | 53 | M | 438.5 | ANT | ANT | aHC, pHC | aHC, pHC | CLB, LTG | Nonlesional | N/A |
| 2 | 23 | F | 368.5 | CM, ANT |  |  | aHC | CLB, LTG, VPA | Nonlesional | N/A |
| 3 | 30 | F | 302 |  | CM, ANT |  | aHC | LTG, OXC | Nonlesional | L |
| 4 | 16 | F | 330.5 | ANT | ANT | aHC | aHC | LTG, LEV, OXC | Nonlesional | L,R |
| 5 | 59 | M | 307 |  | CM, ANT |  | aHC | CBZ, LTG, ZNS | HS, MTS | L |
| 6 | 24 | M | 254 | CM, ANT |  | aHC, pHC |  | LTG, OXC | Nonlesional | R |
| 7 | 22 | F | 214.5 | CM,ANT |  | aHC, pHC |  | CBD, BRV, CNB, ZNS | Left frontal astrocytoma | L |
| 8 | 33 | M | 294 |  | CM,ANT |  | aHC, pHC | CLB, LEV, VPA | Nonlesional | L |
| 9 | 14 | F | 240.5 | CM,ANT |  | pHC |  | LTG | Structural (MCD) | N/A |
| 10 | 42 | F | 216 | ANT, PUL | ANT, PUL | aHC, pHC | aHC, pHC | LEV | Nonlesional | L,R |
| 11 | 26 | M | 278.5 | ANT, PUL | ANT, PUL | aHC, pHC | aHC, pHC | LTG | Nonlesional | L,R |
| 12 | 34 | F | 185 | ANT, PUL |  | pHC |  | CZP, ESL | Structural (PMG/PVNH/SOD) | R |
| 13 | 22 | M | 146.5 | PUL |  | aHC, pHC |  | CLB, OXC | Structural (MTS, FCD) | R |
| 14 | 36 | M | 245 |  | ANT, PUL |  | aHC, pHC | CZP, BRV, OXC | Infectious (encephalitis) | N/A |
| 15 | 65 | M | 186.5 | ANT | ANT | aHC, pHC | aHC, pHC | LTG | Nonlesional | L,R |
| 16 | 39 | F | 220 |  | CM, ANT |  | aHC | LTG, LEV | FCD | L |
| 17 | 31 | F | 269 | ANT | ANT | aHC | aHC | LTG | Nonlesional | L,R |
| 18 | 16 | F | 341 |  | ANT, PUL |  | aHC, pHC | ESL, FFA, RFM, CNB | FCD | L |
| 19 | 42 | F | 409.5 | CM, ANT, PUL | CM, ANT, PUL | aHC, pHC | aHC, pHC | CZP, LEV | SOD, bilateral PVNH | N/A |

THAL = Thalamus, HIPP = Hippocampus, CM = centromedian nucleus, ANT = anterior nucleus, PUL = pulvinar, aHC = anterior hippocampus, pHC = posterior hippocampus, CLB = clobazam, LTG = lamotrigine, LEV = levetiracetam, OXC = oxcarbazepine, ZNS = zonisamide, CBD = cannabidiol, CBZ = carbamazepine, CNB = cenobamate, VPA = valproic acid/depakote, CZP = clonazepam, ESL = eslicarbazepine, FFA = fenfluramine, RFM = rufinamide, MTS = mesial temporal sclerosis, FCD = focal cortical dysplasia, PVNH = periventricular nodular heterotopia, SOD = septo-optic dysplasia, PMG = polymicrogyria, MCD = malformations of cortical development.

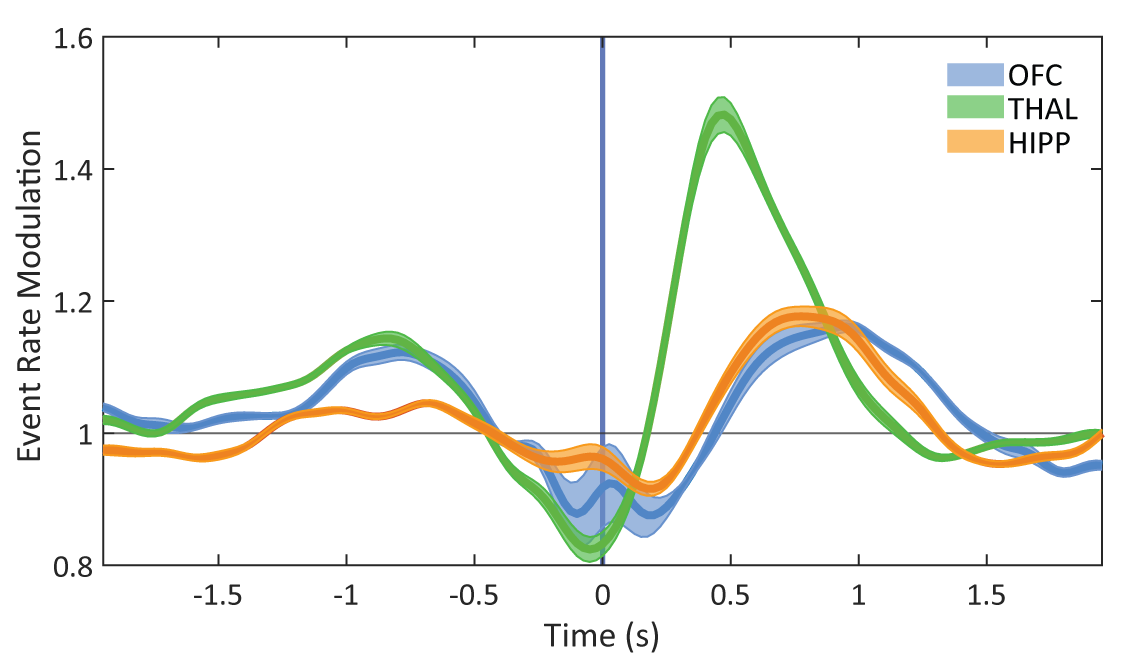

Supplementary Figure 1: Maximum of spindles relative to the OFC down-state. *Modulation in incidence of the maximum amplitude of spindles relative to baseline incidence of spindles in the OFC, thalamus, and hippocampus. Time 0 s marks moment of the OFC slow oscillation minima.*

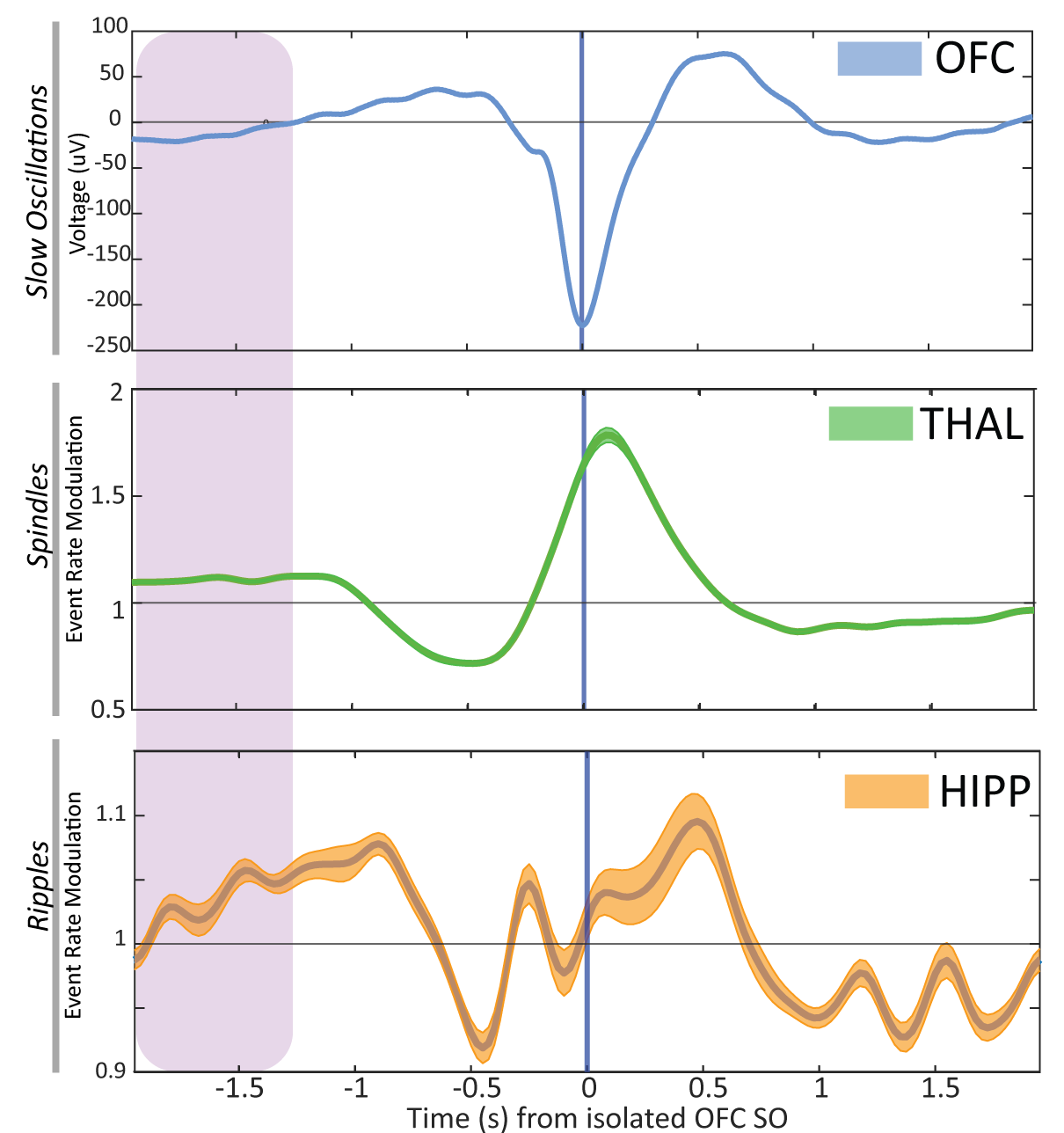

Supplementary Figure 2: Spindle and ripple modulation relative to isolated orbitofrontal cortex slow oscillations. *(top) Modulation in incidence of the thalamic spindles relative to baseline incidence of spindles. (bottom) Modulation in incidence of the hippocampal ripples relative to baseline incidence of ripples. Time 0 s marks moment of the isolated OFC slow oscillation minima.*

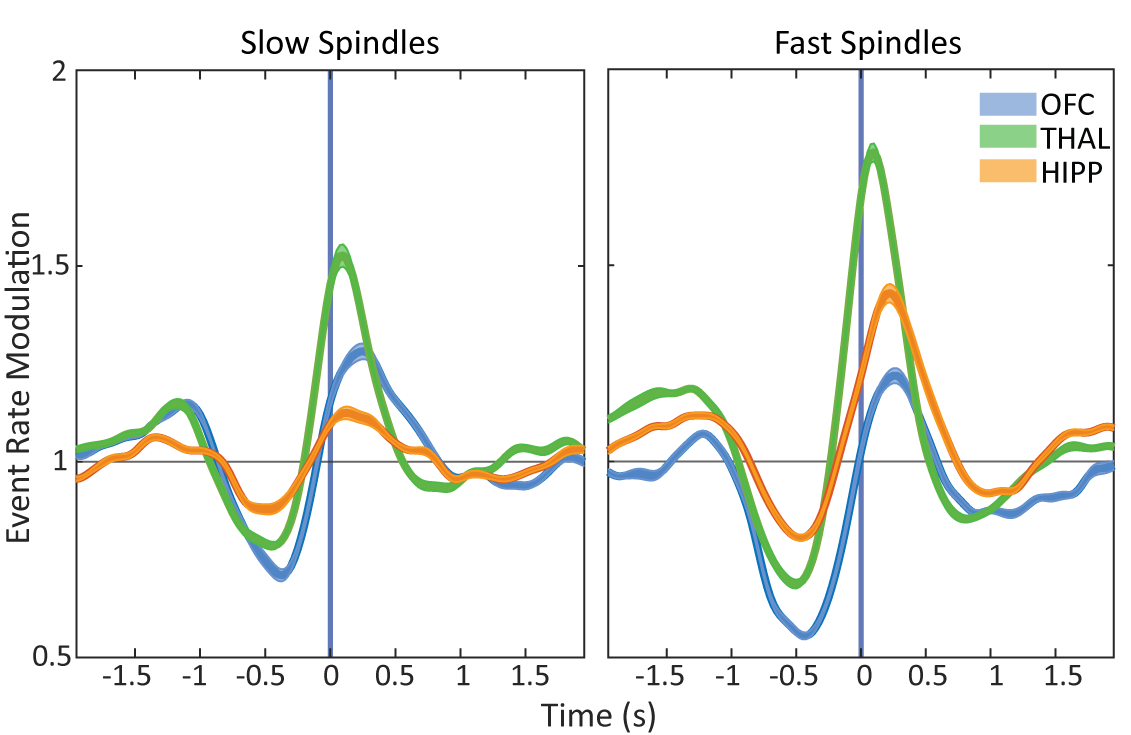

Supplementary Figure 3: Slow and fast spindle probability around OFC slow oscillations. *Modulation in incidence of the onset of slow (left) and fast (right) spindles relative to baseline incidence of (slow/fast) spindles in the OFC, thalamus, and hippocampus. Time 0 s marks moment of the OFC slow oscillation minima.*

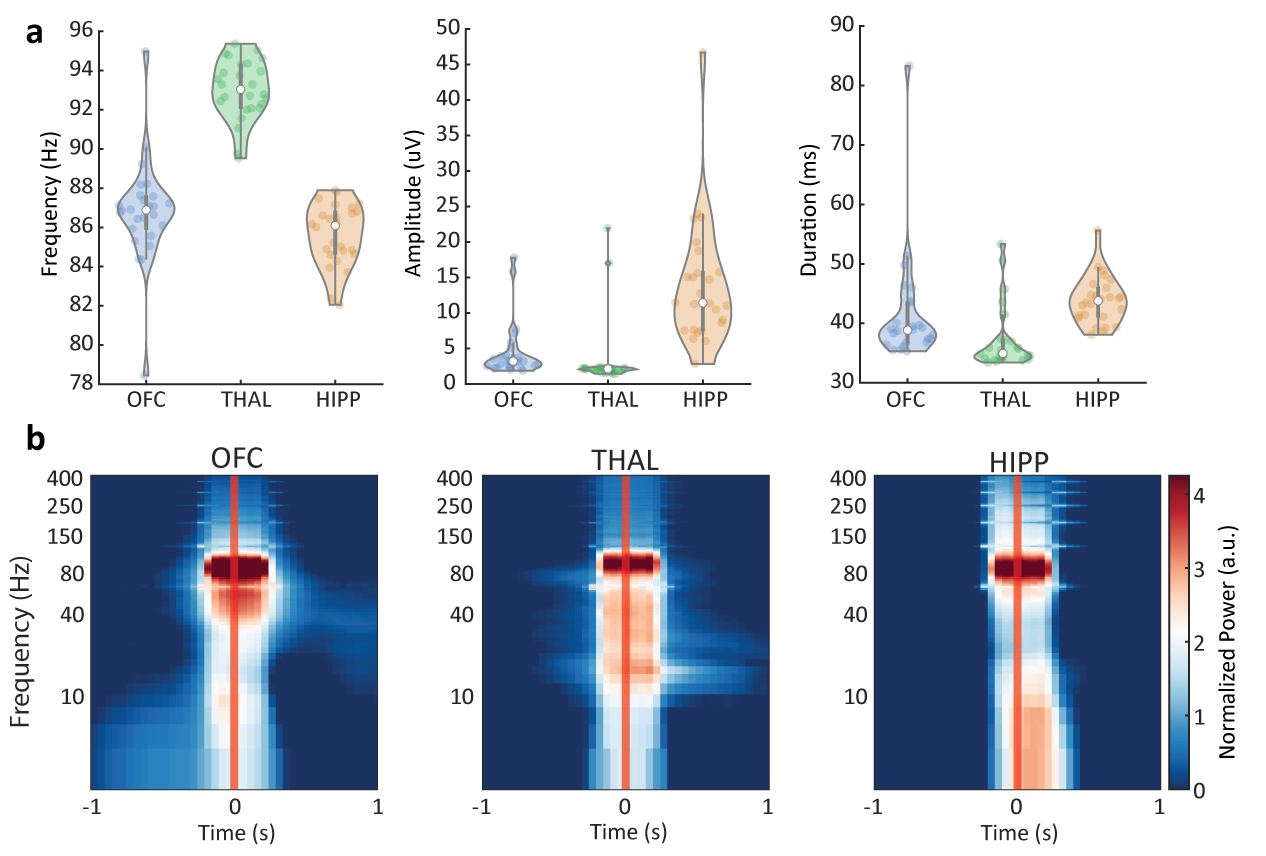

Supplementary Figure 4: Ripple characteristics across the OFC, thalamus and the hippocampus. *(a) Frequency, amplitude and duration of ripples in each region. Every dot represents an individual subject. (b)* *Each column shows the average spectrogram from all subjects representing the evoked average spectral response using a multitaper spectral estimator. Image colors represent the standardized power increase.*

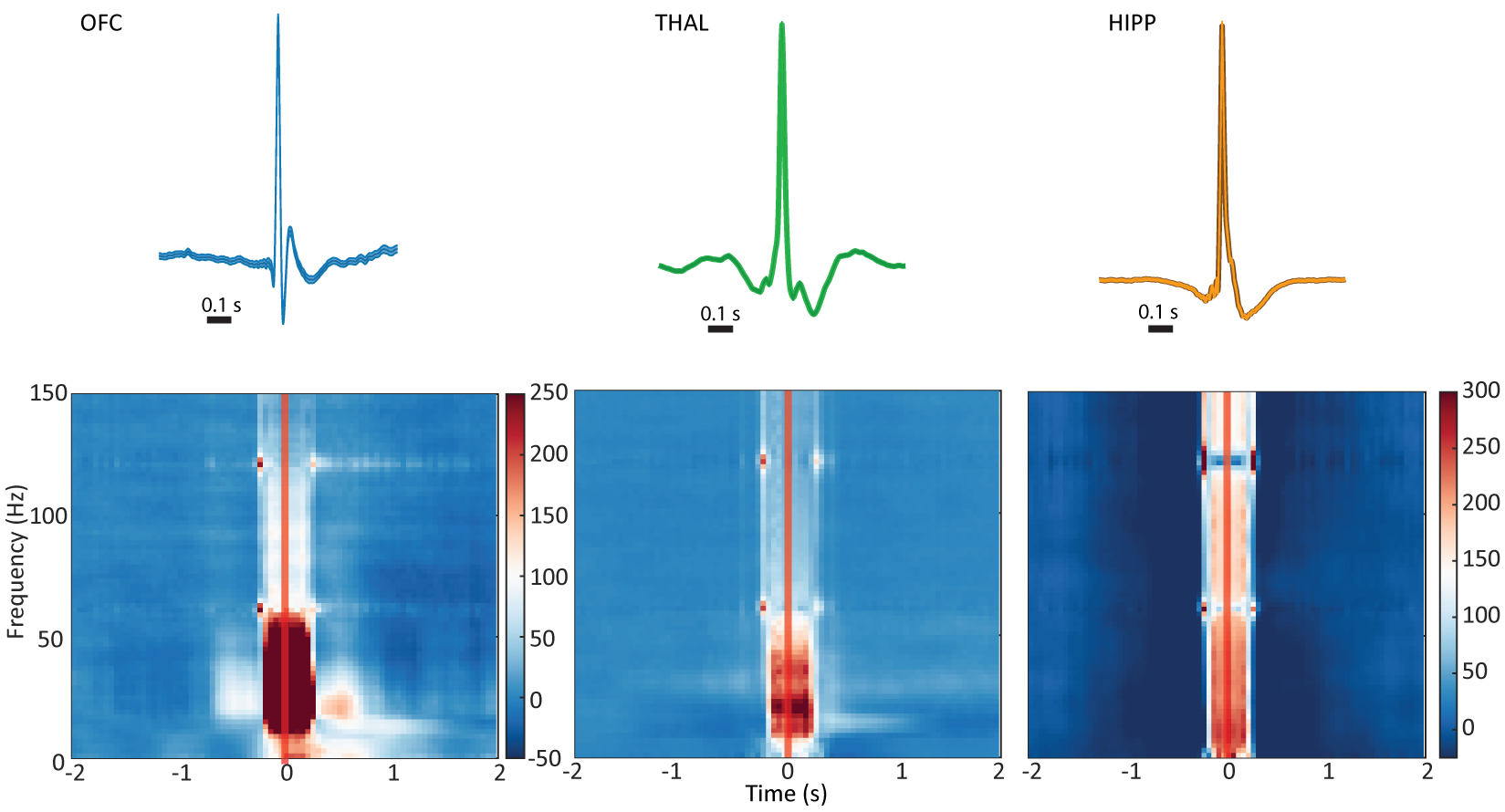

Supplementary Figure 5: Example average epileptic spike waveforms and spectrograms for each region. *Each column shows an average epileptic spike trace from one subject with the spectrogram below representing the evoked average spectral response using a multitaper spectral estimator. Image colors represent the percentage increase in power relative to a baseline interval. Note that the OFC epileptic spike spectrogram is on a different scale.*
